# Genetic Association of *SLC47A1* Gene Variant (17:19571562C >T) and Bioinformatics Analyses of MATE1 Protein in Chronic Kidney Disease Patients of Pakistani Origin

**DOI:** 10.1101/2023.08.12.553072

**Authors:** Mehnaz Ghulam Hussain, Rashid Saif, Amna Younus, Mehmoona Zahid, Sadia Jabeen, Hooria Younas

## Abstract

Chronic Kidney Disease (CKD) is a serious human threat worldwide which is associated with a number of environmental, clinical and genetic factors that affect serum creatinine (SCr) and glomerular filtration rate (eGFR). One of the best biochemical and genetic markers to study renal functioning is *SLC47A1* which encodes MATE1 protein and can be a promising target to study its genetic causes, subject protein is a cationic transporter involved in the regulation of creatinine and urea levels in the blood which may cause toxicity and renal disorders. Hence, the current study focused to interrogate the putative association of *SLC47A1* gene variant (17:19571562C>T) in 75 individuals (cases=50, controls=25) of Pakistani origin using ARMS-PCR genotyping, out of which 10 were found heterozygous, 57 homozygous wild-type and 08 homozygous mutants. PLINK data analysis toolset was used which manifested that sampled population complies the Hardy-Weinberg Equilibrium by χ^2^ (2, *N* = 75) =, *p* = 4.756 × 10^-005^, similarly the Chi-square statistics *p* = 0.03274 along with odds-ratio of 3.244 showing a significant genotypic association with the subject phenotype indicating that mutant allele is almost 3-times more prevalent in cases vs control cohorts with an alternative allele frequency of 0.22 and 0.08 in CKD patients and healthy controls respectively. Besides, few of the bioinformatics tools i.e., ProtParam, PsiPred, PDB-RCSB, Motif Scan, CTU-TMHMM-2.0, ScanProsite, PRmePRed, GPS PAIL2.0, NetOGlyc4.0, NetPhos3, SIFT, Polyphen-2 and STRING were also used for the prediction of different parameters of both the wild-type and mutant MATE1 proteins such as physiochemical properties, secondary structure prediction, 3-D structure, protein conserved domains, transmembrane structures, posttranslational modifications, SNP prediction/amino acid substitution, pathogenicity and protein-protein interaction networks. Current preliminary genotyping provided an insight into the association of the aforementioned genetic variant with CKD to estimate the genetic risks along with prediction of its functional effects in the system biology context which may be considered for developing new targeted drugs along with genetic counselling initiatives to evade this peril in the population.

## Introduction

Chronic Kidney Disease (CKD) is a major global health problem characterized by a progressive loss of kidney function, estimated by the glomerular filtration rate (eGFR) less than 60 mL/min/1.73 m^2^ (1). The probable prevalence of CKD in the general population worldwide is 13.4% ranging from of 11.7%-15.1% (2) although, it is significantly higher in Pakistani population over the age of 50Y with 43.6% and in people under the age of 30Y with10.5% (3). The causes of renal failure are multifaceted and blowout with a combination of genetic, clinical and environmental factors. However, genetic factors are thought to play a substantial role, and several sequence variants have been identified that are associated with an increased risk of renal failure. For instance, the sequence variants of *SLC47A1, RNF128, RNF186, SLC6A19*, and *SLC25A45* have been identified to be associated with an increased serum creatinine level from the normal range of 4.3mg/dL-12.3mg/dL which is an important biomarker of kidney function (4).

*SLC47A1* gene consist of 83,650bp, located on chr.17 p11.2, that encodes MATE1 protein (multidrug and toxin extrusion transporter 1, 570aa), highly expressed at basolateral and apical membranes of renal tubular epithelial cells, liver and muscles in humans (5-7). MATE1 protein shows homology with toxic compound extrusion transporter NorM in bacteria, which worked by the electrochemical gradient of protons and cations across the cell membranes (6, 8-12). The MATE1 protein also plays an important role for the secretion/extrusion of creatinine, metformin, cimetidine and tetraethyl ammonium (TEA) into the urine (13).

In the *SLC47A1* gene, more than 70 polymorphisms have been identified as “Stop-gain mutations” and “frameshift INDELs” that cause the loss of gene functioning (4,14) however, in the current study, rs111653425 p.(Ala465Val) variant identified as missense mutation in *SLC47A1*, that alters the amino acids composition of transmembrane domains within the long cytoplasmic loop of renal tubule and causes the dysfunction of MATE1 transporter, lead to the elevation of SCr, eventually resulted in CKD and renal damage (6, 11). Creatinine secretion through MATE1 protein can be interfered by certain drugs e.g., cimetidine, trimethoprim, corticosteroids, pyrimethamine, phenacemide and salicylates (15).

The current research was designed to evaluate genetic association of *SLC47A1* gene variant with CKD patients of Pakistani origin. The subject gene is located at Chr.17, accession ID (NC_000017.11) in assembly GRCH38.P14 (GCF_000001405.40). This variant is located at 17:19571562C>T position with dbSNP ID (rs111653425) in exon 15^th^ of the gene according to GenBank transcript accession (M55158.1) at its r.1480 and CDS position of (c.1394C>T) along with NP_060712.2: (p.Ala465Val) protein locus.

## Materials and Methods

### Sample collection and DNA extraction

A total 75 samples were collected including 50 adult CKD patients attending the nephrology outpatient department of the Akhtar Saeed Trust Hospital, Life Care Hospital and Farooq Hospital (Lahore-Pakistan) between July-August 2023 along with 25 healthy controls to study the association of *SLC47A1* gene variant g.17:19571562C>T with CKD. The criteria for selection were age bracket (19-65) years, urea level range (71-217mg/dL) and serum creatinine level (4.3-12.3mg/dL). The patient’s blood was taken into the EDTA-K3 vacutainers and stored at 4°C (1, 2) Genomic DNA were extracted from collected samples using GDSBio (https://www.gdsbio.com/en/) kit by following the manufacturer guidelines (3). Samples details are provided in Supplementary Table (S1).

### Primer designing

MATE1 proteins complete CDS was downloaded from GenBank accession ID: NC_000017.11 and ARMS-PCR primers were designed manually by OligoCalc & NetPrimer softwares. In total five primers were designed, three of which are “reverse normal (N) ARMS”, “reverse mutant (M) ARMS” and a “forward common” for amplification of normal and mutant alleles. A secondary mismatch was also added at the fourth nucleotide position from the 3’-end of the reverse wild/mutant ARMS primers to improve the effectiveness and specificity of the primers. To amplify a region near the target sequence, two additional primers (forward and reverse) were designed that serve as an internal control (IC) (4). Details of primers are given in (Table 1).

**Table 1:**
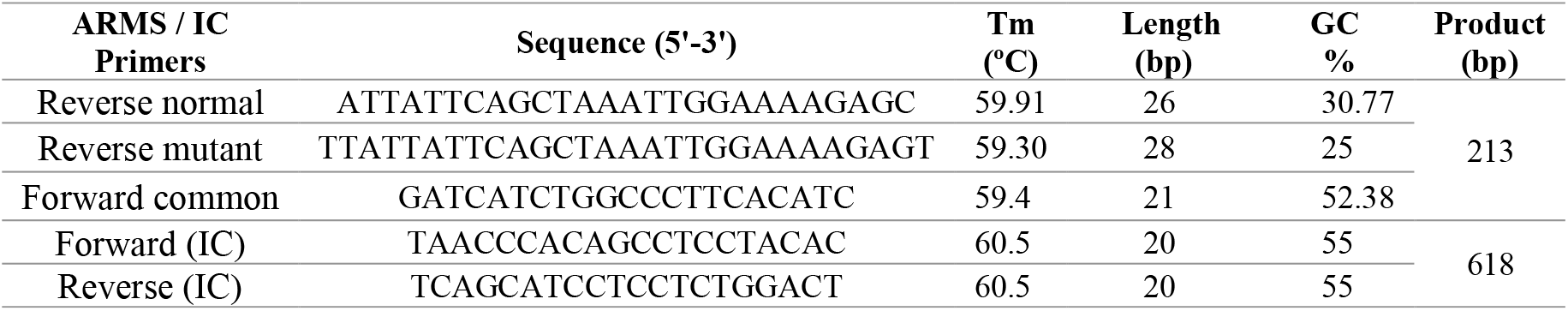
ARMS-PCR primers sequences and other attributes.

### DNA amplification

SimpliAmp thermal cycler (Applied Biosystem) was used following the ARMS genotyping technique. Analysis of each sample was done in two separate PCR reactions having common forward with mutant-ARMS reverse primer in one tube and common forward with normal-ARMS reverse primer in the second tube along with IC primer-set were used in both tubes to check the fidelity of PCR amplification. In this protocol of PCR 2.5mM dNTPs, 0.051U/μL *Taq* polymerase, 2.5mM MgCl_2_, 2uL of extracted 50ng/μL of genomic DNA, 10mM of each primer, 1x *Taq* buffer and DEPC treated molecular grade water was used in a total of 16uL reaction mixture. The PCR protocol was adopted with 5-minutes of initial denaturation at 95°C followed by 30 cycles of denaturation at (95°C for 30 sec), annealing at (60°C for 30 sec), extension at (72°C for 30 sec) with the final extension of 72°C for 10 minutes and finally stored at 4°C (Figure 1) (5, 6).

**Figure 1:**
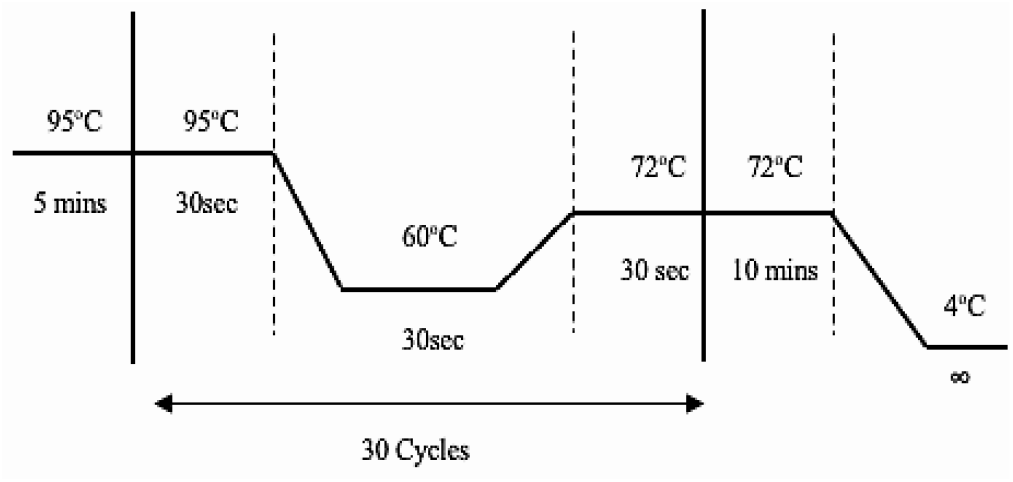
Thermal cyclic conditions for ARMS-PCR

### Statistical analysis

To compute the observed and expected genotypic and allelic frequencies, Hardy Weinberg Equilibrium (*HWE*) was used by obeying *p*2+2*pq*+*q*2 = 1 equation and Chi-square analysis using 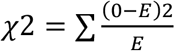 to check the association of subject variant (rs111653425C>T) with the creatinine level in sampled population using PLINK data analysis toolset. Additionally, allele frequencies in cases and controls were also calculated along with association *p*-value and odds-ratio (7, 8). Besides, SPSS bivariate analysis was further performed to assess the correlation between creatinine and urea levels (9).

### Bioinformatics analysis

To check the different properties of wild and mutant-type MATE1 proteins in the present study *in-silico* approach was adopted and a number of bioinformatic tools/databases were utilized. The transcript of the MATE1 protein was obtained from NCBI and its mutant sequence was obtained by incorporating the subject DNA/amino acid variant and translated through EMBL-EBI. The BLAST was run to check the similarity/homology between wild and mutant protein sequences and it was observed that both sequences are the same except for an amino acid change of (A>V) at 465 position.

### Physiochemical properties

“ProtParam” tool (http://web.expasy.org/protparam/) was used to analyze the physiochemical properties, atomic composition, theoretical PI, instability, amino acids composition, molecular weight, instability, aliphatic index, extinction coefficient, GRAVY and estimated half-life in both mutant and wild-type protein sequences of MATE1 (10). To achieved this, FASTA sequence of both wild and mutant-type proteins were uploaded and the results were concluded.

### Secondary structure prediction

To predict the secondary structure of wild and mutant-type protein “PsiPred” (http://bioinf.cs.ucl.ac.uk/psipred/) tool was used that predicts alpha helices, beta sheets and coils in protein structures. For this the FASTA files of both proteins were submitted with default parameter and Visual Molecular Dynamics (VMD) was used to view the structure of predicted proteins (11).

### Three-dimensional (3D) protein structure prediction

To explore the 3D structure of MATE1 wild and mutant-type protein, “PDB RCSB” (https://www.rcsb.org/) tool/database was utilized (12). The FASTA sequences of both proteins were uploaded one by one and their structures were analyzed and compared.

### Protein conserved domain prediction

To evaluate the conserved regions in the wild and mutant-type MATE1 proteins “Motif scan” tool was used. At first, protein sequences of both proteins were pasted in search bar, option for “motifs” was selected and launched the search (13).

### Transmembrane structures estimations

“CTU-TMHMM” server v.2 (https://services.healthtech.dtu.dk/service.php?TMHMM-2.0) was used to predict the transmembrane helices, intracellular and extracellular regions in the wild and mutant-type MATE1 proteins by using FASTA sequences.

### Post-translational modifications assessments

The post translational modifications are important for protein functioning. In the current research, these modifications in wild and mutant-type MATE1 protein were also predicted with the help of number of available tools such as “ScanProsite” (http://prosite.expasy.org/scanprosite/) tool for phosphorylation, “GPS_PAIL2.0” (http://bdmpail.biocuckoo.org/prediction.php) tool for acetylation, “PRmePRed” (http://bioinfo.icgeb.res.in/PRmePRed/#) tool for methylation and “NetOGlyc-4.0” (https://services.healthtech.dtu.dk/service.php?NetNGlyc-1.0) tool for glycosylation (15-18).

### Valuation of impact of variants on protein function

The Sortin Intolerant from Tolerant (SIFT) tool was used for measuring the impact of non-synonymous SNPs and the biological pathogenicity of *SLC47A1* variant (rs111653425) for wild and mutant-type MATE1 proteins. A mutation is estimated as deleterious if the score is <0.05 (14).

### Estimation of effects of amino acid changes on protein functions

In the present study, Polymorphism Phenotyping v2 (PolyPhen-2) tool was used to predict the impact of an amino acid substitution on the structure as well as function of wild and mutant-type MATE1 proteins using straightforward physical and comparative considerations (15).

### Protein-protein interactions illustration

STRING is an online bioinformatics tool/database that illustrates the protein-protein interactions (16). To evaluate the protein interactions (direct and indirect) of wild and mutant-type MATE1 proteins, the sequence was run on tool and generated results were analyzed.

## Results

### Variant Genotyping

In the current study, *SLC47A1* variant (rs111653425) in Pakistani population showed the significant association with CKD, that is in concordance to the other population reports in the world (17) and depicts the under-selection nature of this Chr.17 locus 19571562 C>T in large human population. A total of 75 samples (cases=50, controls=25) were genotyped (Figure 2). Statistical analysis revealed that out of total genotyped cases, 08 were heterozygous, 36 homozygous wild type and 6 homozygous mutants, whereas in controls, 4 samples were heterozygous and 21 homozygous wild-type while none of the sample was observed as homozygous mutant. All inclusive, 10(13%) heterozygous, 57(76%) homozygous wild-type and 8(10%) homozygous-mutants were found in the sampled population. Furthermore, the OR calculated as 3.244, which indicates that almost 3 times mutant allele is more prevalent in cases than controls.

**Figure 2:**
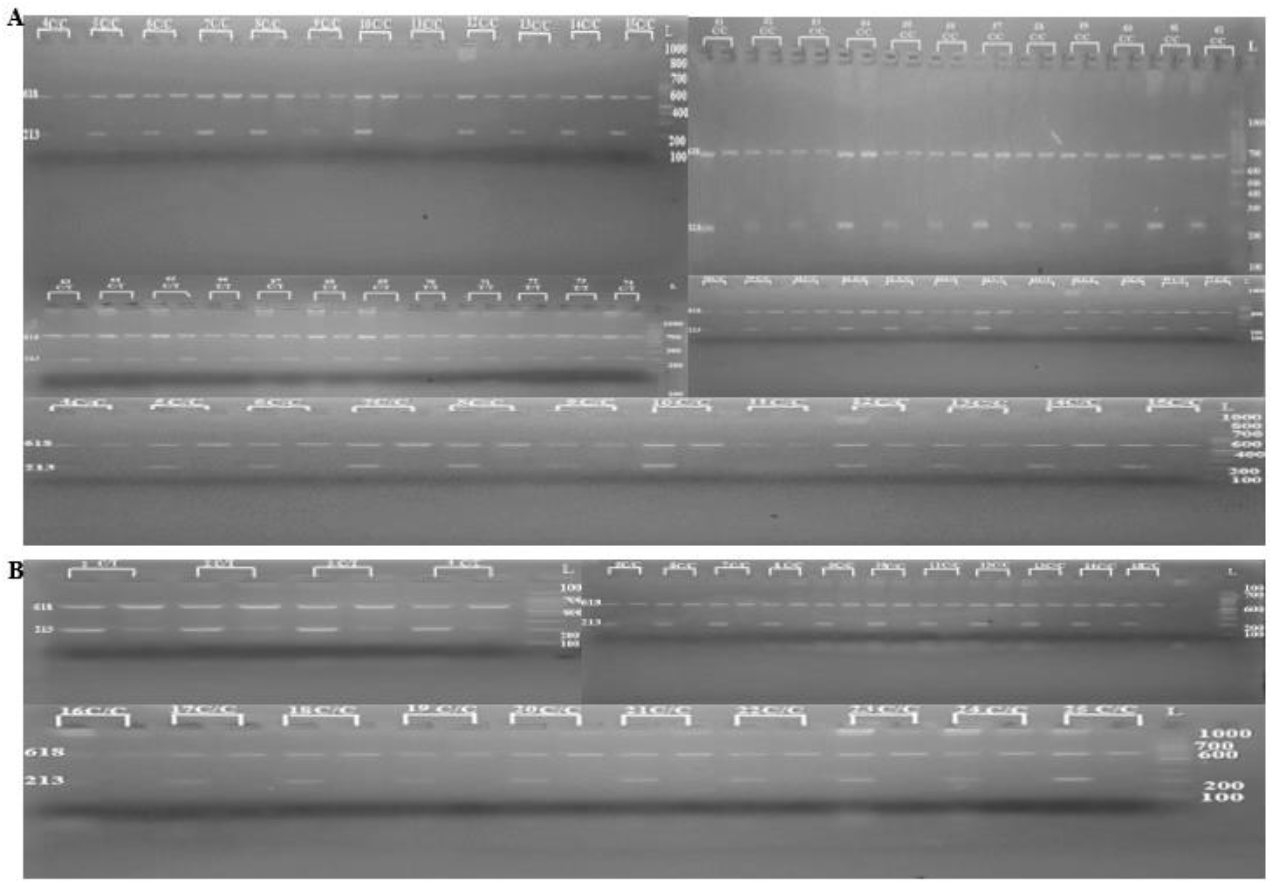
Genotyping of 50 CKD cases (A) and 25 controls (B)

### Statistical association/correlation analysis

To check whether our sampled population is obeying the principle of Hardy Weinberg Equilibrium (HWE) or not, Chi square analysis was conducted with the following outcomes (*p* = 4.756 x 10^− 005^), which rejecting our null hypothesis as the *p*-value is below the set threshold confidence interval of 0.05.

Furthermore, using PLINK data analysis toolset genetic association was conducted which showed that alternative alleles frequencies of 0.22 and 0.08 were observed in cases and controls respectively having significant *p-*value = 0.03274 and OR of 3.244, signifying a considerable association of the screened variant with CKD. Furthermore, above mentioned OR displaying the prevalence of odds/mutant is almost 3 times higher in cases as compare to controls (Table 2).

**Table 2:** Association analysis of subject variant in Pakistani human population.

| # of Samples | Chr. Position | cDNA variant NM_018242.3 | Protein variant NP_060712.2 | Genotypic information | | | Alternative allele frequency | | $p$-value (OR) |
| --- | --- | --- | --- | --- | --- | --- | --- | --- | --- |
|  |  |  |  | Homo-wild (CC/%) | Hetero- (CT/%) | Homo-mutant (TT/%) | Cases | Controls |  |
| 75 | 17:19571562 | c.1394C>T | NP_060712.2 | 57/76 | 10/13 | 8/10 | 0.22 | 0.08 | 0.03274 (3.244) |

Eventually, Pearson correlation test was applied with SPSS software to check correlation between three variables of age, urea and creatinine. It was analyzed that there is week-to-moderate positive correlation of urea and creatinine was predicted with 0.435, week negative correlation was observed in age and creatinine level, similarly, very week positive correlation was observed in age and urea levels. Additionally, Chi-square analysis showed highly significant correlation prediction among aforementioned variables with each other as the observed *p*-value is 0.0001 (Table 3).

**Table 3:** Pearson correlation analysis of age, creatinine and urea levels.

| Variables | Correlation | p-value |
| --- | --- | --- |
| Urea/creatinine | .435 | 0.0001 |
| Age/creatinine | -0.117 |  |
| Age/urea | .099 |  |

### SLC47A1 protein analysis using different bioinformatics tools

#### I. Physiochemical findings of MATE1 by ProtParam

ProtParam tool was utilized to analyze physiochemical properties of MATE1 proteins (both wild and mutant-type) encoded by *SLC47A1* gene. Statistically, it was found that proteins have 570aa with molecular weight 61922.18 Daltons, and their theoretical isoelectric point (PI) was 7.53. The chemical formula for both proteins was C_2844_H_4531_N_727_O_763_S_26_ but their atomic numbers were different which were 8885 for wild-type and 8891 for mutant protein. However, aliphatic index, instability index and Grand average of hydropathicity (GRAVY) of both proteins were also variable. The observed value forming cysteine was 67225 and reduced value was 66350 in both protein variants. Similarly, positively charged residues (Arg+Lys) were 35, whereas the negatively charged (Asp+Glu) residues were 34. The half-life of mammalian protein under in vitro conditions was estimated as 30 hrs while in vivo it was predicted to be more than 10. (Table 4).

**Table 4:** Physicochemical properties of wild and mutant type proteins.

| Physicochemical Properties | (Wild-type) | (Mutant-type) |
| --- | --- | --- |
| Amino acids | 570Aa | 570Aa |
| Molecular weight | 61922.18 | 61922.18 |
| Theoretical PI | 7.53 | 7.53 |
| Molecular formula | C<sub>2842</sub>H<sub>4527</sub>N<sub>727</sub>O<sub>763</sub>S<sub>26</sub> | C<sub>2844</sub>H<sub>4531</sub>N<sub>727</sub>O<sub>763</sub>S<sub>26</sub> |
| Number of atoms | 8885 | 8891 |
| Negative charged residues (Asp + Glu) | 34 | 34 |
| Positive charged residues (Arg + Lys) | 35 | 35 |
| Extinction coefficient | 67225(assuming all pairs of Cys form cysteine)<br>66350(assuming all pairs of Cys are reduced) | 67225(assuming all pairs of Cys form cysteine)<br>66350(assuming all pairs of Cys are reduced) |
| Estimated half life | 30hrs | 30hrs |
| Instability index | 37.06 | 36.14 |
| Aliphatic index | 126.04 | 126.32 |
| GRAVY | 0.646 | 0.650 |

The detailed analysis of protein sequence, revealed that the amino acid Leucine (L) was present in high numbers with 93(16.3%), Valine (V) was 56 (9.8%), Alanine (A) and Glycine (G) were 49(8.6%) and 48 (8.4%) while the other amino acids were observed between 0(0%) to 37(6.5%) (Figure 3A). These numbers of amino acids are also confirmed by graphical analysis presented in (Figure 3B), the amino acid Leucine is showing high peak of concentration. There is no variation in concentrations of both wild-type and mutant MATE1 proteins.

**Figure 3:**
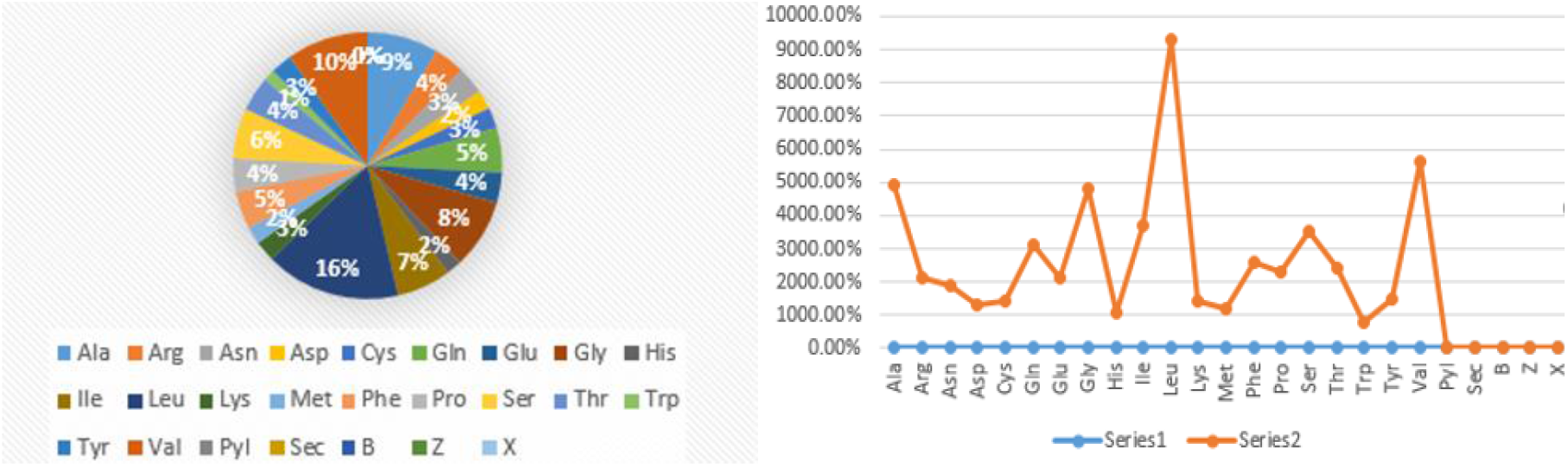
Atomic ratios of MATE1 **(A)**, amino acid percentages **(B)**

#### II. Secondary structure predictions of MATE1 using PsiPred

Secondary structures including helices, beta strands and coils of both wild and mutant-type proteins were analyzed using PsiPred tool. The pink shaded amino acids are showing helices and light grey color indicates coils. The difference in both proteins is at amino acid number 465 which is Val (V) in place of Ala (A) in mutant protein (Figure 4).

**Figure 4:**
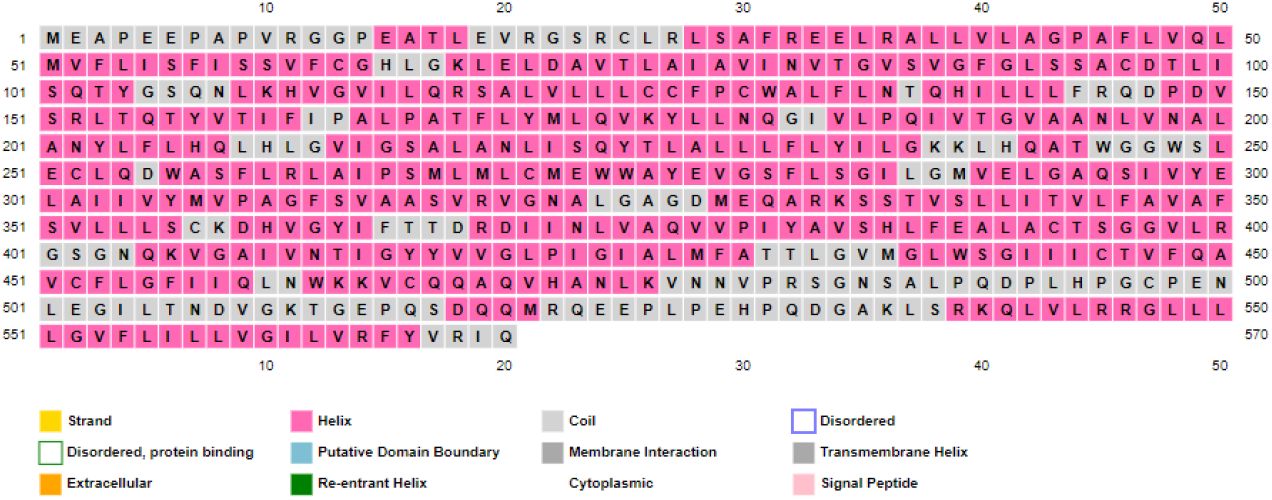
Amino acids involved in making secondary structures

#### III. 3D structure prediction of MATE1 using PDB RCSB

PDB RCSB tool determined 3D structure of both (wild and mutant-type) MATE1 proteins showing carbohydrates and inner chains polymer structures (Figure 5). However, no change in 3D structures of both wild and mutant-type proteins was observed.

**Figure 5:**
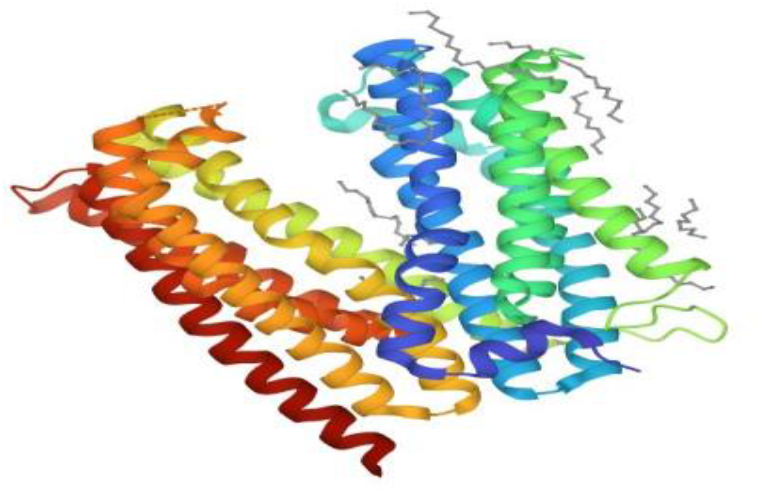
3D structure of MATE1protein

#### IV. Protein domain predictions using Motif Scan

Motif Scan tool predicted number of motifs present in MATE1 protein and results showed only one motif is present in both wild-type and mutant proteins with pfam ID (MatE), two positions along with E-values are 44…204 (8.6e-31) and 265…424 (3.2e-31) respectively (Table 5).

**Table 5:** Domain prediction in MATE1.

| pfam ID | Position (independent E-value) | Description |
| --- | --- | --- |
| MatE | 44..204(8.6e-31)<br>265...424(3.2e-31) | PF01554, MatE |

#### V. Transmembrane helices (TMHs) finding using TMHMM tool

According to TMHMM analysis, 13 TMHs are present in 570 amino acids long MATE1 protein, out of which 280 amino acids are present in these TMHs. The rest of amino acids are present on inner and outside of TMHs. No difference in number of helices in both wild and mutant-type MATE1 proteins was observed. According to graph the purple line indicates transmembrane portion of amino acids, blue line shows presence of amino acids inside the helices and the orange line represents amino acids outside the TMHs in both proteins (Figure 6).

**Figure 6:**
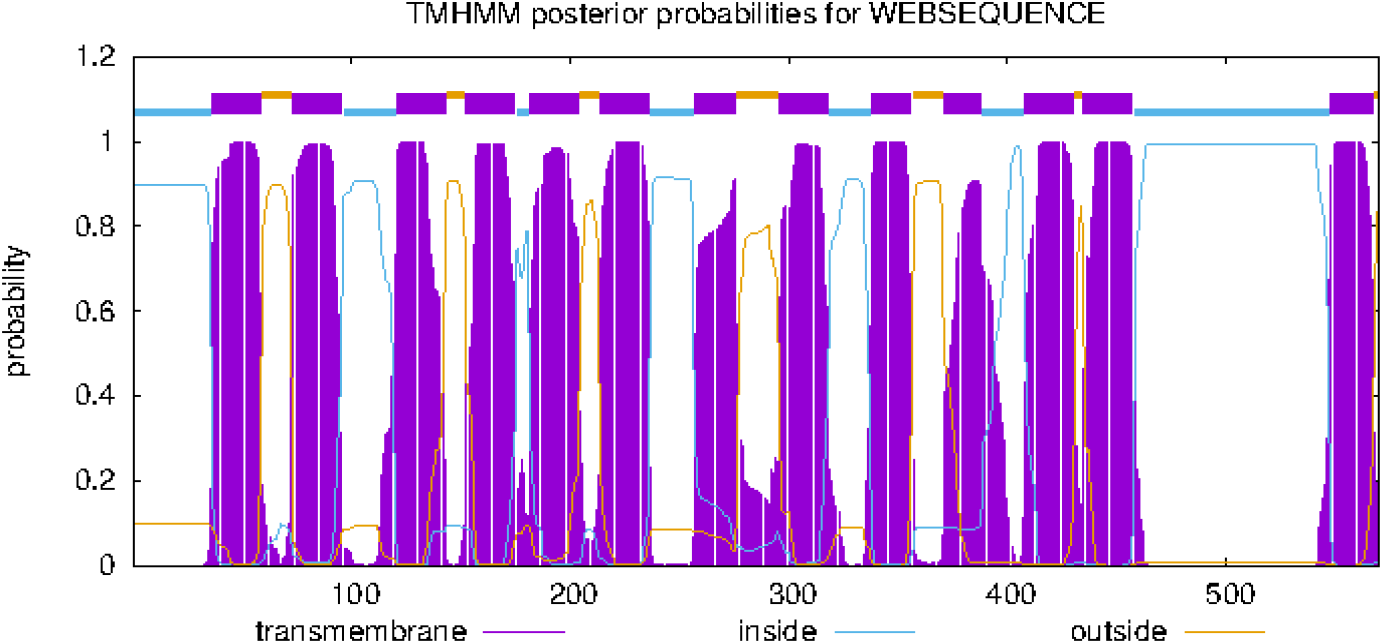
Transmembrane helices (TMHs) in MATE1 protein

#### VI. Post-translational modifications findings using ScanProsite

The occurrence of patterns, profiles and motifs in proteins (wild and mutant-type) was analyzed by ScanProsite tool. Additionally, detailed understanding of post-translational parameters such as glycosylation, phosphorylation and cell attachment sequences were also computed by this tool. Vaults are large ribonucleoprotein particles having a main component (MVP) which is made up of 53 amino acids. In MATE1 protein only one MVP domain is present between 484-545 amino acids and its score is 4.930 (Figure 7).

**Figure 7:**
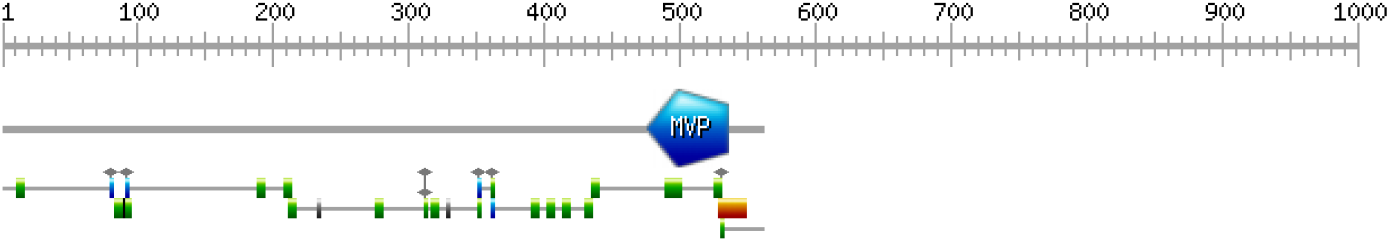
Post-translational modifications prediction in MATE1 protein

##### a. Glycosylation and phosphorylation findings of the MATE1 residues

Determination of C-linked, N-linked, O-linked or S-linked glycosylation sites by identity of amino acid atom which covalently binds the chain of carbohydrates. In MATE1 protein the N-linked glycosylation predicted site is 82 and phosphorylation is at serine (S) and threonine (T) amino acids. In MATE1 protein serine phosphorylation is at 94, 317, 356, 538 whereas at threonine it is at position 367 (Table 6).

**Table 6:** Glycosylation and phosphorylation sites of MATE1 protein.

| Amino acids positions | Glycosylation | Phosphorylation |
| --- | --- | --- |
| 82 | N-linked (GlnNAc...) | ----- |
| 94 | ----- | Phosphoserine |
| 317 | ----- | Phosphoserine |
| 356 | ----- | Phosphoserine |
| 367 | ----- | Phosphothreonine |
| 538 | ----- | Phosphoserine |

Leucine zipper sequence (LsrkqlvLrrlllLgvflilL) is in-between 537-558 amino acids N-Myristoylation (Table 7).

**Table 7:**
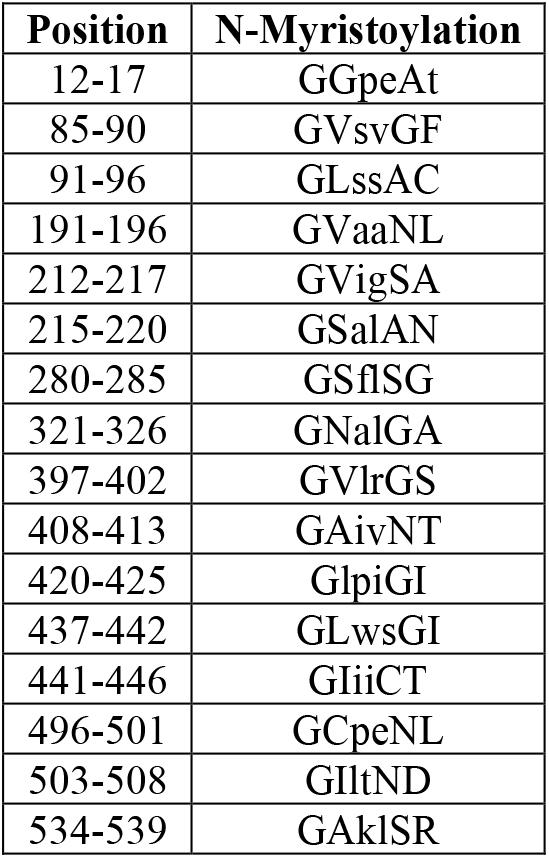
Leucine zipper sequence.

##### b. R and K residues of methylation of MATE1 by PRmePRed

The PRmePRed showed the predictions for all R and K residues of methylation of MATE1 (wild and mutant-type) proteins which were predicted at 11, 21, 24, 27, 32, 36, 319, 333, 400, 482, 539, 545, 546 positions with different peptide sequences and prediction scores respectively (Table 5).

**Table 5:**
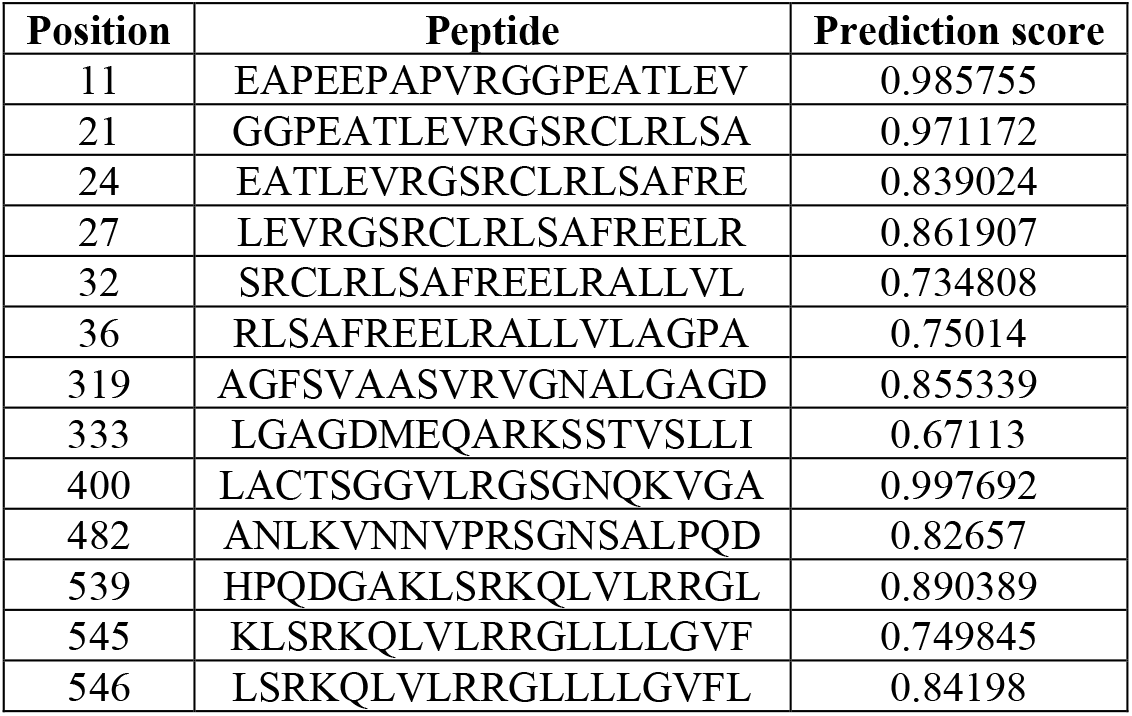
Methylation sites of MATE1.

##### c. Glycosylation predictions of MATE1 using NetOGlyc-4.0

The NetOGlyc-4.0 tool showed glycosylation on carbohydrates which bind to targeted MATE1 protein. Purple line in the graph represents the threshold value 0.5 which ranges from (0-1) and threshold value above 0.5 of any residual indicates the greater possibility of glycosylation while the green line shows potential for glycosylation from (0-500) and the graph is same for both wild and mutant-type MATE1 proteins, no difference to predict glycosylation possibility was observed in both proteins (Figure 8).

**Figure 8:**
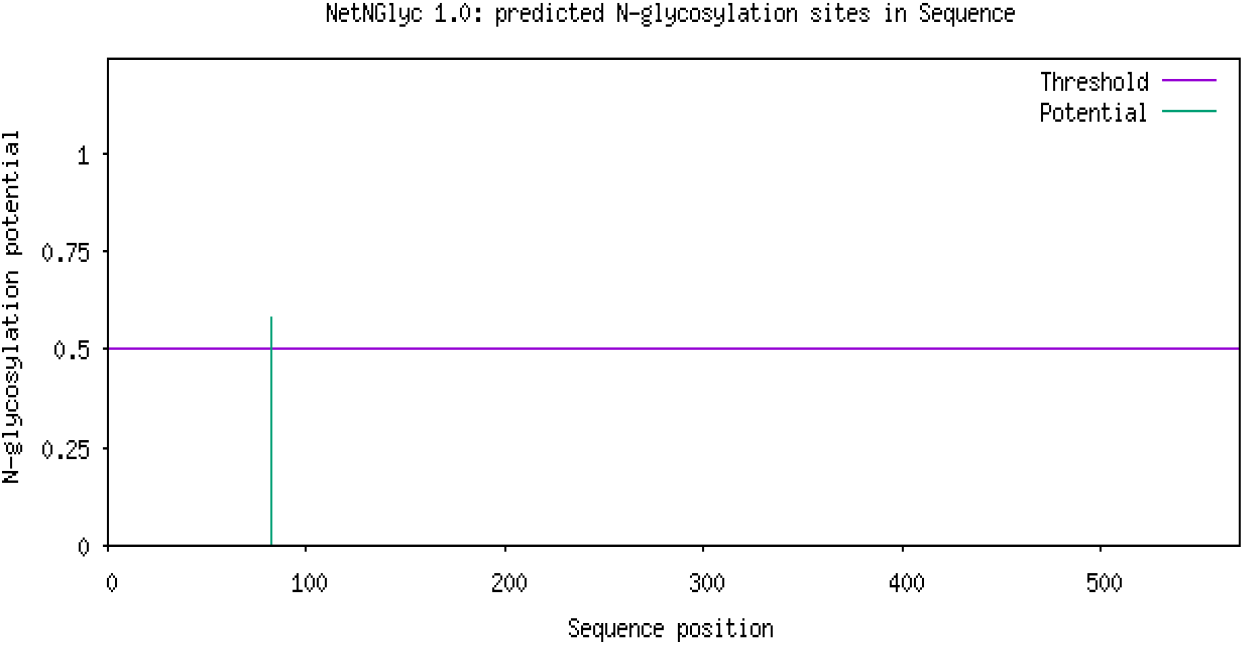
Glycosylation sites of MATE1 protein

##### d. Acetylation of lysine (K) prediction in MATE1 by GPS PAIL 2.0

Acetylation of lysine (K) residues in the MATE1 (wild and mutant-type) proteins sequence is at the position K536 and KAT2A distribution sites are 100% predicted by the GPS PAIL 2.0. No difference in acetylation sites of lysine (K) in both wild and mutant-type MATE1proteins was observed (Figure 9).

**Figure 9:**
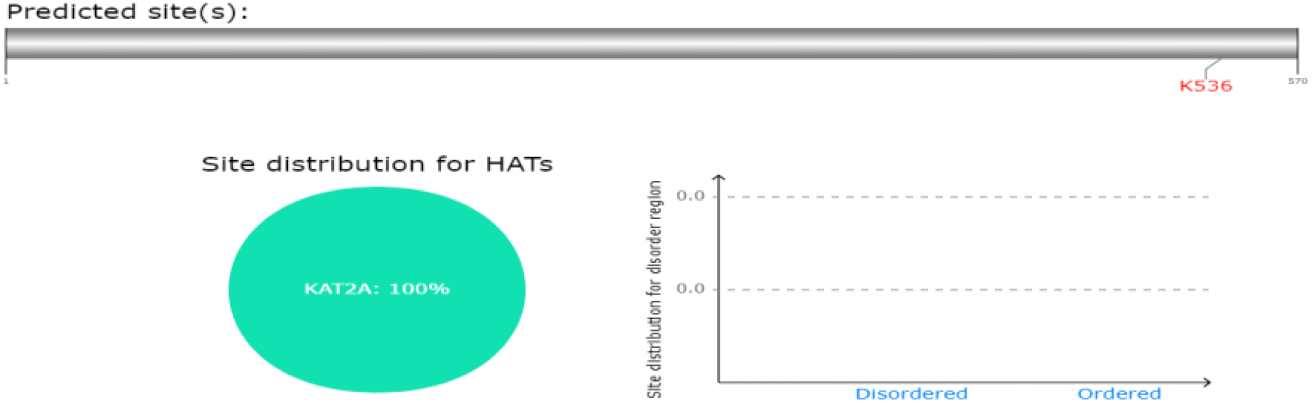
Acetylation of lysine (K) in both MATE1 variant

##### e. Tyrosine, serine and threonine phosphorylation sites prediction by NetPhos 3.1

NetPhos 3.1 analyzed tyrosine, serine and threonine phosphorylation sites in the sequence of both wild and mutant MATE1 proteins. According to (Figure 10) threshold limit is indicated by pink line which is (0-1) but for MATE1 protein the observed threshold value is 0.5 and higher threshold value indicates more possibility of phosphorylation. Blue line shows phosphorylation on tyrosine, green line on threonine and red line on serine respectively. However, no difference in phosphorylation sites was observed in both wild and mutant-type proteins.

**Figure 10:**
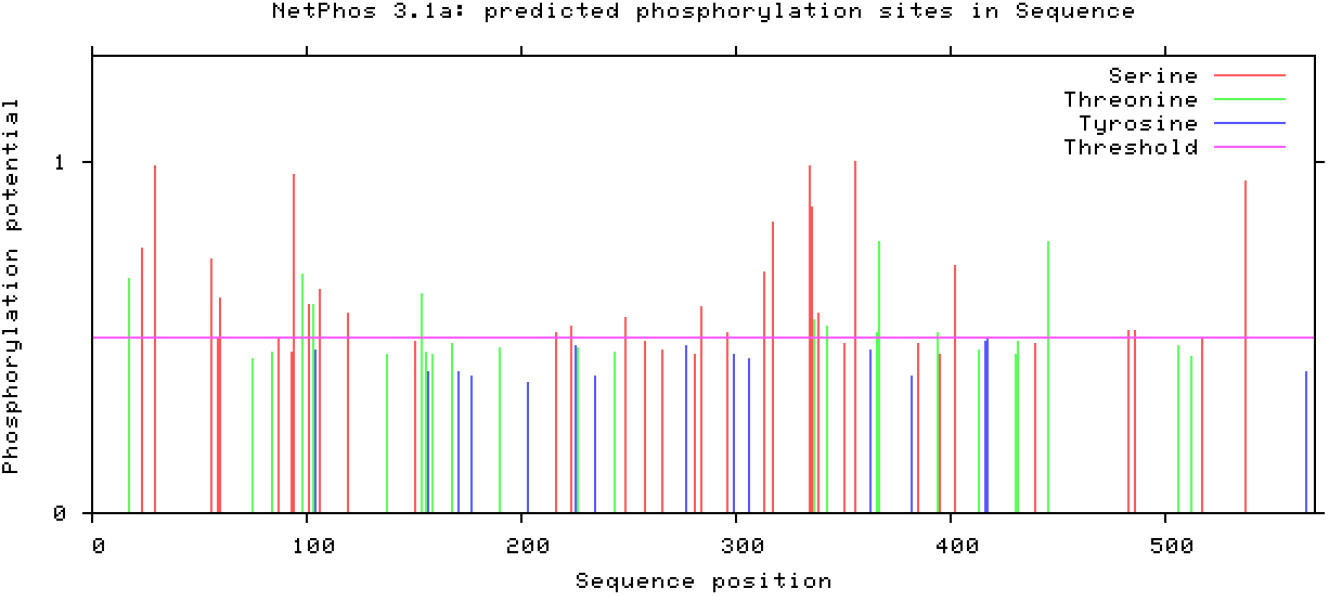
Tyrosine, serine & threonine phosphorylation sites in MATE1 protein variants

#### VII. Nonsynonymous SNP predictions using SIFT

SIFT tool analyzed nonsynonymous missense mutations (SNPs) in the MATE1 protein and the amino acids mutation was identified in mutant MATE1 protein at position 465 in place of Ala (A) the amino acid Val (V) is present and its threshold is 0.91 which is more than the set threshold 0.05, it means it is deleterious but predicted as tolerated. Black color shows nonpolar amino acids, green indicates uncharged, red shows basic and blue color shows acidic amino acids. The substitution at position 465 predicts to affect protein function with a score of 0.04, median sequence 3.36 which is higher than 3.25 thus showing a warning (Figure 11).

**Figure 11:**
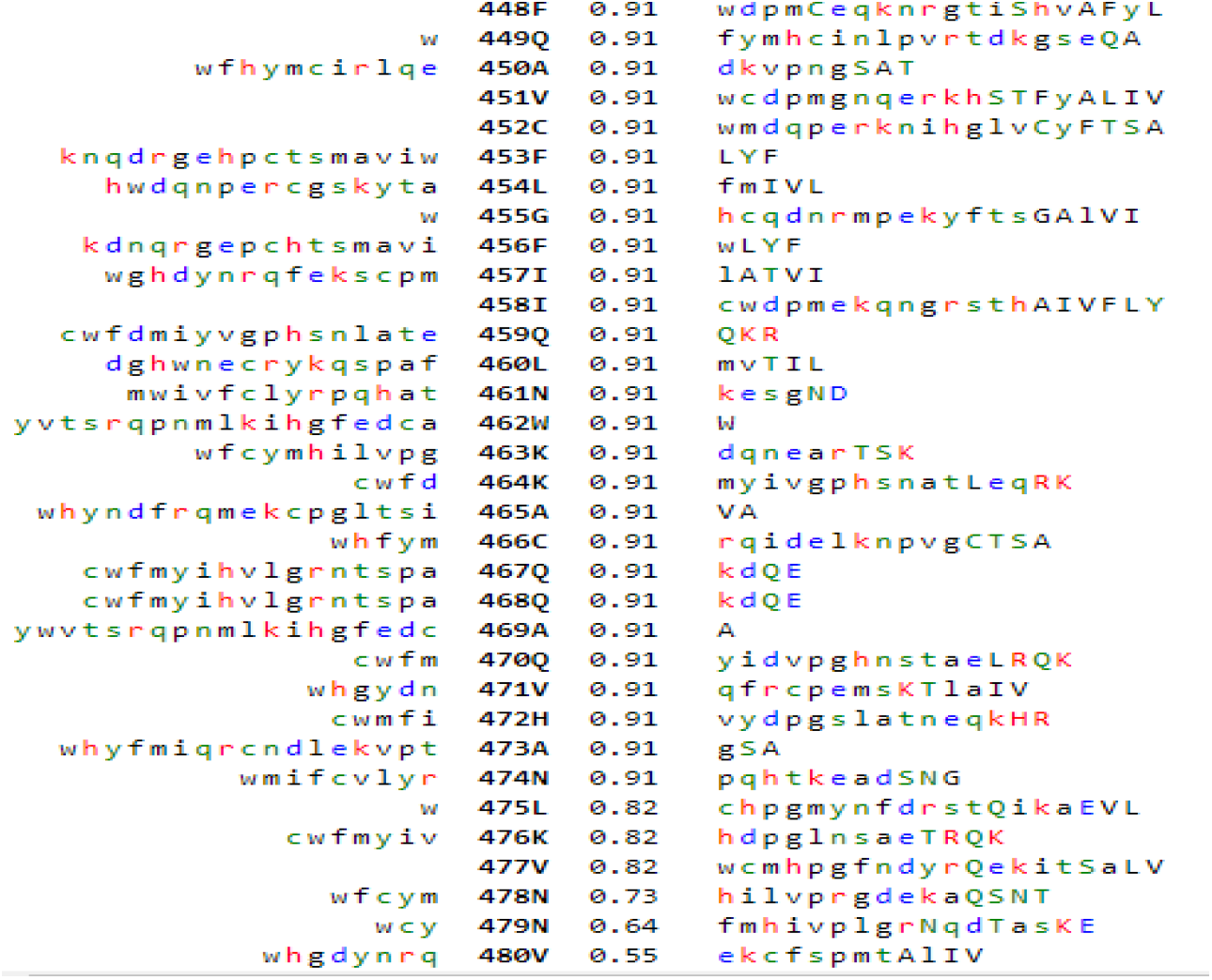
Nonsynonymous SNP prediction in mutant-type MATE1 protein

#### VIII. Pathogenicity depictions of MATE1 by Polyphen2

The Polyphen-2 tool diagnosed that in mutant MATE1 protein the mutation is ‘BENIGN’ with a prediction score of 0.100 (sensitivity:0.93) and (specificity:0.85) (Figure 12).

**Figure 12:**
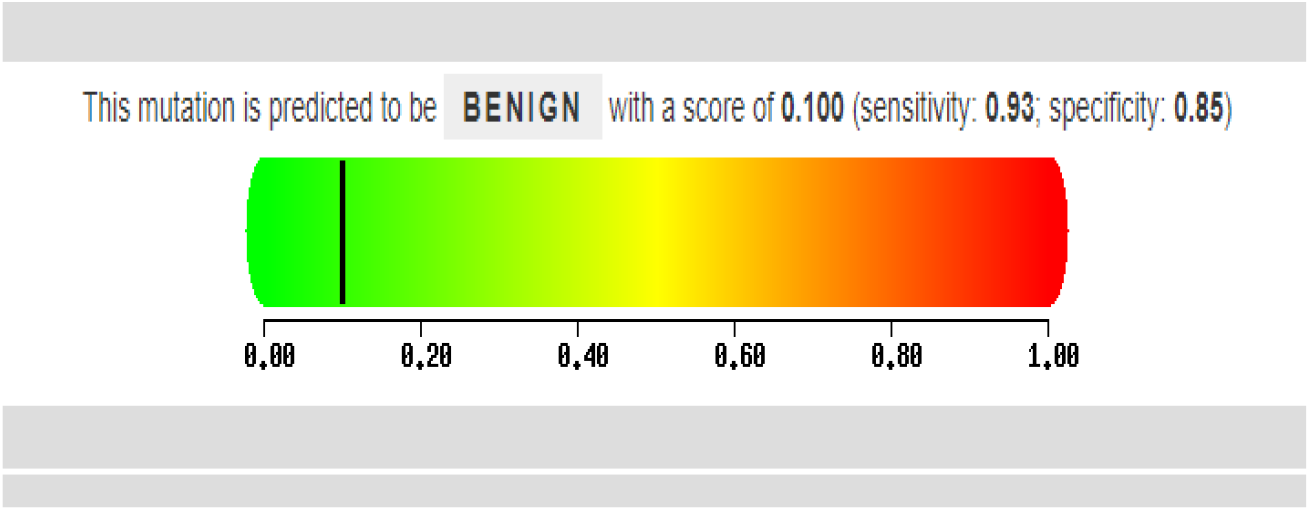
Pathogenicity depiction of mutant-type MATE1 protein

#### IX. Protein-protein interactions using STRING database

STRING analysis showed interactions of *SLC47A1* (MATE1) protein with other SLCs proteins and it depicted the interaction of *SLC47A1* with other genes such as *SLC29A4, SLC22A4, SLC22A8, SLC01B1* etc. as shown in (Figure 13).

**Figure 13:**
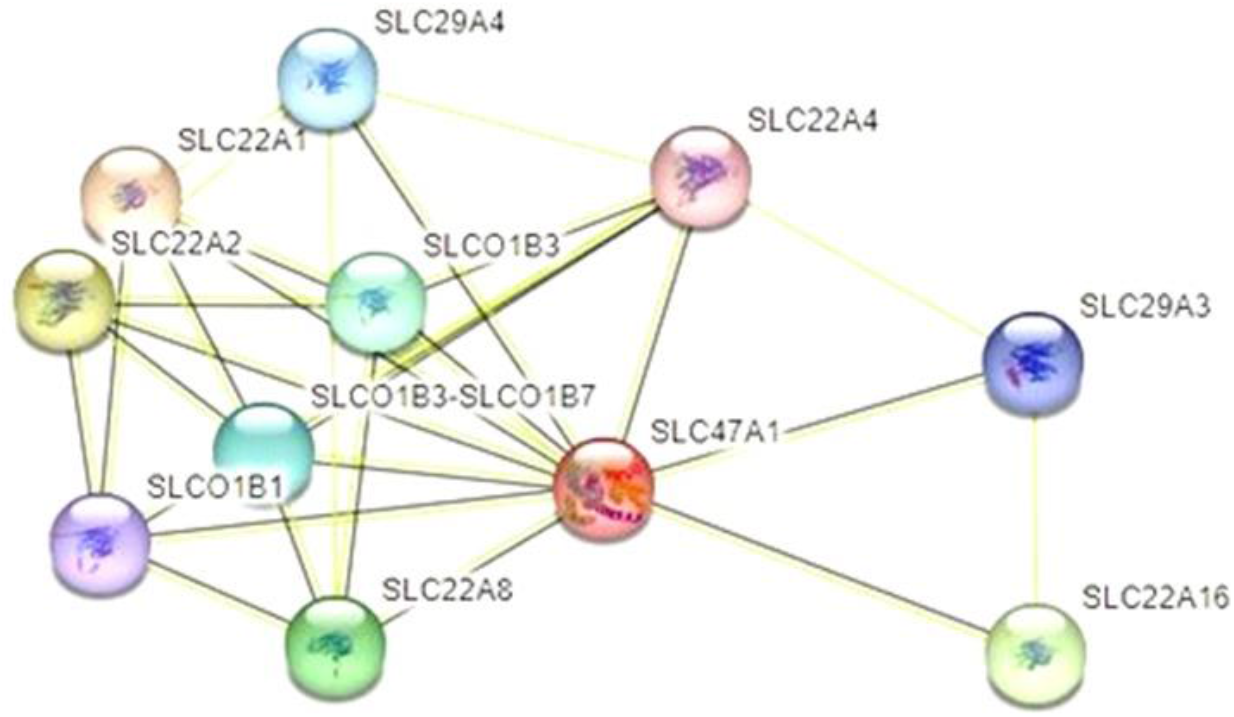
String analysis of protein-protein interactions of MATE1 protein

Around 11 nodes are present connecting solute-carriers family proteins, nodes represent proteins and shells of interaction. Colored nodes represent query proteins and first shell of interactors, white nodes show second shell of interactors, empty nodes show proteins of unknown 3D structure and filled nodes represent 3D structure is known or predicted. Similarly, edges represent protein-protein interactions in which the blue edge represents known interactions curated by the database while purple edge shows experimentally determined known interactions. The green line shows the gene neighborhood in predicted interactions, red lines show gene fusions and blue lines show gene co-occurrence. Similarly other interactions where yellow edge represents text mining interactions, black are the co-expressions and sky blue are protein homology.

In addition, following statistics were also observed that there are 28 edges, average node degree is 5.09, average local clustering coefficient is 0.473 and PPI enrichment *p*-value is 2.93×10^-06^ for both wild and mutant type MATE1 proteins as shown in (Table 6).

**Table 6:** Protein-Protein Interactions of *SLC47A1* (MATE1) protein.

|  |  |
| --- | --- |
| Number of nodes | 11 |
| Number of edges | 28 |
| Average node degree | 5.09 |
| Avg. local clustering coefficient | 0.473 |
| Expected number of edges | 10 |
| PPI enrichment p-value | 2.93e-06 |

## Discussion

The kidneys are the vital and multifunctional organs, filter metabolic wastes and transport essential substances, such as electrolytes and water by glomerular filtration in the glomerulus, a small ball of capillaries in the kidney. The tubular part of the nephron, reabsorbs essential substances and secretes waste products into the tubule lumen, made up of 14 distinct components, each of which performs specific transport processes, mediated by transporter genes, ion channels, ATPase and solute carriers (18).

In exceptional cases, both kidneys may stop working, which can be a life-threatening condition, but many people with CKD are able to put up active lives with proper treatment and health-conscious lifestyle, such as nutritious diet and workout. Genome-wide association studies have shown that in the *SLC47* family, MATE1 (SLC47A1) protein is an organic cation/proton exchanger, mostly expressed in renal tubular cells and hepatocytes, can transport creatinine and drugs such as metformin, paraquat or the anticancer drug e.g., oxaliplatin (19).

MATE1 was identified in 2005, as a mammalian homologue of the bacterial NorM protein, an H+/organic cation antiporter, which means that it transports organic cations out of cells in exchange for protons. This suggests that MATE1 also mediates the electroneutral exchange of organic cations in renal excretion and significantly transports creatinine, metformin, cimetidine, and tetraethyl ammonium (TEA) into the urine. These are all organic cations that are either naturally produced by the body or taken as medications. The regulatory mechanisms that control the expression of the MATE1 gene were studied with a focus on the basal level of transcription. A deletion analysis, mutational analysis and electrophoretic mobility assay were used to identify Sp1 as a general transcription factor for human MATE1, that binds to the GC-rich region of the MATE1 promoter. This binding recruit TATA-binding proteins and fixes the transcription start site which is essential for the basal level of expression of the MATE1 gene (20) (Figure 13).

**Figure 13:**
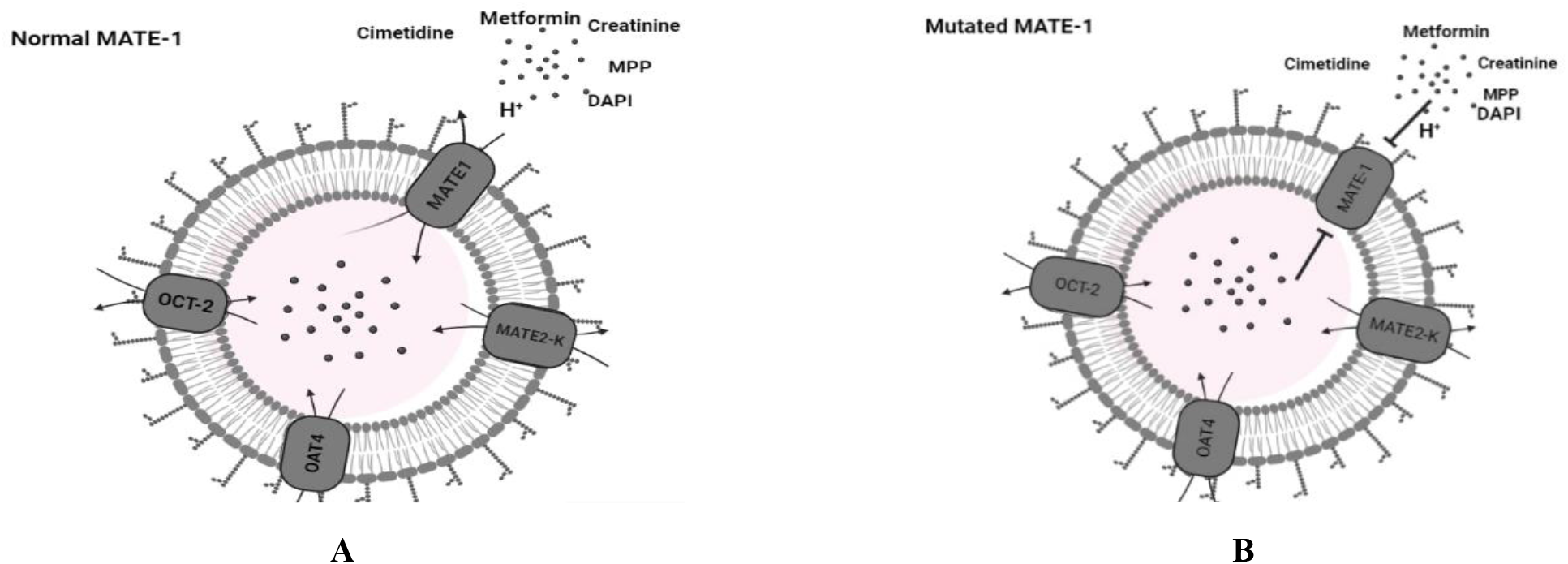
Pathophysiology of normal mechanism of MATE1 transporter **(A)**, a mutated protein mechanism **(B)**

Any mutation in *SLC47A1* resulted in the malfunctioning of MATE1 protein that produced accumulation of creatinine (a substrate for MATE1). In gene *SLC47A1* more than 70 polymorphisms have been identified “Stop-gain mutations” and “frameshift INDELs” have the largest impact of sequence variants to loss of function of a gene. However, in the current study, rs111653425 variant identified as missense mutation p.(Ala465Val) in *SLC47A1*, that results in different amino acid being incorporated into the structure of a MATE1 protein resides transmembrane domains within the long cytoplasmic loop of renal tubule, and alter the function of MATE1 transporter, eventually leads to the elevation of SCr and resulted in CKD or renal damage (6, 11). Creatinine secretion through MATE1 protein can be affected by certain drugs (cimetidine, trimethoprim, corticosteroids, pyrimethamine, phenacemide and salicylates) (15).

Previously, it was highlighted in number of studies that mutation in locus 17:19571562C>T is significantly associated in developing CKD. It is a complex disorder with a genetic component (21) and was found that rare variants in three solute carriers *SLC6A19, SLC25A45* and *SLC47A1* and two E3 ubiquitin ligases (RNF186 and RNF128) are associated with serum creatinine (SCr) in Icelanders. These variants have a larger effect on SCr than previously reported common variants and they may affect kidney function or creatinine synthesis and excretion. Three of the variants were also associated with CKD (21).

The present study aimed to find association of variant 17:19571562C >T (rs111653425) with CKD in Pakistani population and found enough evidence of association of the subject variant with development of CKD in Pakistan. It was observed that out of total 75 sampled population, 10 were heterozygous, 57 were homozygous wild-type and 08 were homozygous mutants. Hence, it can be concluded that mutant allele is almost 3-times more in cases vs control cohorts with an alternative allele frequency of 0.22 and 0.08 in cases and controls respectively in Pakistani population at the highly conserved locus of 17:19571562 (Figure 14)

**Figure 14:**
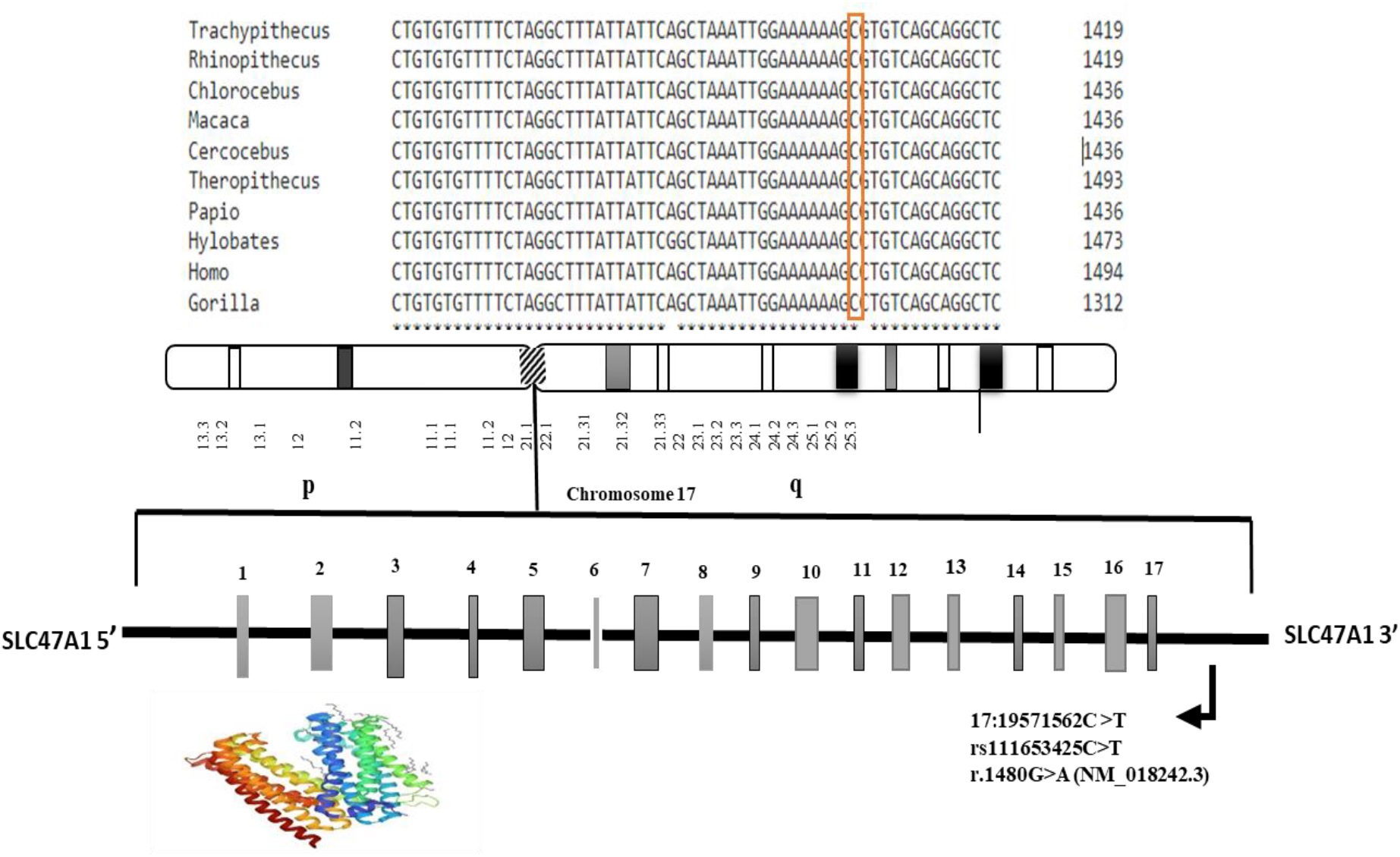
MSA of genotyped locus in ten species along with genetic map and structure of the MATE1 protein

Moreover, the structural and functional features of MATE1 protein is obtained by bioinformatics tools in the current study. Using ProtParam tool chemical and physical parameters of both wild and mutant-type protein (MATE1) were predicted. Secondary structures including helix, beta strands and coils of both wild and mutant-type proteins were also studied using PsiPred tool.

The difference in both proteins is in amino acid number 465 which is V in place of A in mutant protein. The presence of 13 transmembrane helices (TMHs) in this 570 residuals protein was analyzed by TMHMM tool and these TMHs contain around 280.59874 amino acids. Remaining amino acids are on inner and outside of TMHs.

ScanProsite tool was applied to analyze the presence of profiles, motifs and patterns. Furthermore, post-translational parameters like phosphorylation, glycosylation and sequences of cell attachment were also studied which are important for functional specialization in the cell. MATE1 protein contains only one MVP domain between 484-545 amino acids and its score is 4.930. The N-linked glycosylation site is 82 in MATE1 protein. Prediction of phosphorylation was at serine (S) and threonine (T) amino acids. In MATE1 protein serine phosphorylation is at 94, 356, 317, 358 whereas at threonine it is at position 367. Between 537-558 amino acids there is Leucine zipper sequence (LsrkqlvLrrlllLgvflilL).

In brief, we have found a rare protein-coding variant that found associated with elevated SCr in CKD patients from Pakistani origin as previously reported in Iceland (21). Subsequent in-silico analyses revealed that mutated protein is not altering much of the metabolic pathways and intriguing disease conditions/pathogenicity but further in-vivo experimentations are needed to strengthen our understanding about the functional aspects of the subject variant.

## Conclusion

In this study the *SLC47A1* gene variant (rs111653425C>T) was found associated with CKD in selected Pakistani population as the *p* = 0.03274 with OR= 0.344 and alternative allele frequency of 0.22 and 0.08 in cases and controls. Further screening of the subject variant in a larger population would reinforce our initial understandings about this biomarker to assess the risk of onset of CKD and would also help to adopt preventive measures to minimize the risk of this disease through genetic counselling initiatives.

## Acknowledgement

Authors are obliged to Mr. Ahmad Saim for his help in sample collection.

